# The neurodevelopment of delay discounting for monetary rewards in pre-adolescent children

**DOI:** 10.1101/2020.08.10.245316

**Authors:** Mei Yu, Tongran Liu, Jiannong Shi

## Abstract

Children are found to exhibit high degrees of delay discounting compared with adults in many delay discounting studies. However, the temporal dynamics of the differences behind the behaviors are not well known. In this study, we chose two age groups of participants and adopted event-related potential (ERP) techniques to investigate the neural dynamic differences between children and adults during delay discounting processes. Behavioral findings showed that children discounted more than adults and chose more immediate choices. Electrophysiological findings revealed that children exhibited longer neural processing (longer P2 latency) than adults during the early detection and identification phase. Children showed less cognitive control (smaller N2 amplitude) than adults over the middle frontal areas, and they devoted more neural effort (larger P3 amplitudes) to making final choices than adults. The factors of reward amount and time delay could influence the development of delay discounting in male children.

## Introduction

Pre-adolescent children are often characterized as impulsive decision makers and frequently ignore the long-term benefit of their choices. When making a choice between a smaller and sooner option and a larger and longer option, children are more likely to be driven by reward immediacy, while adolescents are more likely to be driven by reward amount (Scheres et al., 2006). It can be assumed that adults may also be driven mainly by reward amount, and children may choose smaller and sooner alternatives more than adults (Steinberg, 2009). For example, when facing the choice of receiving 10 dollars now or obtaining 20 dollars after a month, more children than adults will choose the option of receiving 10 dollars now. The result of differently weighting the two options leads to the subjective value of 10 dollars now being higher than 20 dollars after a month, even though the objective value is inverse. This kind of choice, referring to the decrease in a reward’s subjective value as the delay to its receipt increases, is called delay discounting (Ainslie, 1975; Chapman and Elstein, 1995; Frederick et al., 2002; Green et al., 1996; Green et al., 2004; Richards, 1999; Odum, 2011; Robertson and Rasmussen, 2018; Scheres et al., 2013).

In the widely used paradigm of studying delay discounting (Mitchell, 1999), the immediate choices varied in terms of the amount (from $0.01 to $10.50) received immediately, whereas the delayed choices varied in terms of the time delay (from 0 days to 365 days), but the amount was fixed at $10. The immediate choices and delayed choices were paired in advance but were presented to participants randomly. The delay discounting phenomenon is often described by the hyperbolic model from Mazur (1987), which is written as:

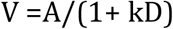

where V denotes the subjective value of an outcome, A denotes its objective value, D denotes the delay to its receipt, and k denotes the rate of discounting (Mazur, 1987). The discounting rate k means the rate at which the subjective value of a reward decreases in value with respect to the time it takes to receive it (Myerson and Green, 1995). A larger k value indicates that the person’s degree of discounting for the delayed reward increases more rapidly (Richards et al., 1997), and the person is more likely to choose immediate choices. Mitchell et al. ‘s (1999) paradigm was widely used in children (Costa Dias et al., 2013; Wilson et al., 2011) and adults (de Wit et al., 2007; Lockenhoff et al., 2011), and its adaption was used in event-related potential (ERP) (Gui et al., 2016) and functional magnetic resonance imaging (fMRI) methods (de Water et al., 2017). Moreover, it has been found that children’s discounting rate was higher than that of adults and decreased with age in many studies (Green et al., 1994 [children aged around 12 years old]; Prencipe et al., 2011 [childhood between 8 and 12 years old]; Scheres et al., 2014 [children of 6 to 12 years old with mean age about 9 years old]; Steinberg, 2009 [children between 10 and 12 years old]). However, the neural mechanisms for the neurodevelopment of delay discounting processes in children have been less studied.

Prefrontal and parietal areas are believed to support higher cognitive functions activated uniformly by delay discounting choices irrespective of the delay, as well as greater relative fronto-parietal activity when subjects chose longer-term options (Ballard and Knutson, 2009; de Water et al., 2017; Figner et al., 2010; McClure et al., 2004; Sripada et al., 2011). These brain areas are also important for the neurodevelopment of top-down systems and delay discounting processes (Blakemore and Choudhury, 2006; Casey et al., 2005; Giedd et al., 1999; Gogtay et al., 2004; Klingberg et al., 2002; Peters and Buchel, 2011; Fuster, 2002; Steinbeis et al., 2016). Christakou et al. (2011), testing a sample of individuals between 12 and 31 years old, found a linear decrease in discounting with age and discovered that the ventromedial prefrontal cortex (VMPFC) contributed to the development of self-control by connecting delay-related information from brain areas such as the dorsolateral prefrontal cortex (DLPFC), insula, and parietal cortex. Moreover, it was found that 12- and 16 year-old adolescents showed a close connection between delay sensitivity and the activation of cognitive control areas and a close relationship between amount sensitivity and the activation of reward valuation areas (de Water et al., 2017).

Several ERP components are also related to delay discounting processes. The frontal P2 component, which occurs approximately 150-200 ms poststimulus onset, is a good index for the early detection and identification of task-related perceptual representations (Potts et al., 2006). It was discovered that P2 amplitudes varied as the time delay increased from 2 weeks to 50 years (He et al., 2012). P2 responses were larger with long delays than with short delays and larger with large delays than with small delays (Gui et al., 2016). It was also found that impulsive adults showed delayed P2 responses (Wu et al., 2016) and that P2 amplitudes increased and latencies decreased with age development in children during an oddball task (Tsai et al., 2012).

Also, the frontal N2 component is referred to in decision making, as it often appears between 200 and 350 ms after stimulus onset and is involved in cognitive control processes, such as cognitive flexibility and inhibitory control (Bruin, 2002; Espinet et al., 2012; Folstein and Van Petten, 2008). Neural activations in cognitive control areas were associated with delayed choices, and larger N2 amplitudes were related to nonimpulsive choices; thus, N2 may act as a neural marker for one’s ability to resist immediate rewards in delay discounting (Gui et al., 2016; McClure et al., 2004). Children aged 7 years old with more activation of the DLPFC showed more negative N2 amplitudes, indicating that more cognitive control resources were recruited (Lamm et al., 2014).

In addition, the parietal P3 is a positive and large-amplitude ERP component with a typical peak latency between 300 and 400 ms over central-parietal areas (Nieuwenhuis et al., 2005). This component reflects the response to stimulus evaluation and decision making (Nieuwenhuis et al., 2005) and is also thought to be related to the processing capacity and mental workload (Kok, 2001). During delay discounting processes, P3 amplitudes were significantly larger in short-term delays (involving more nonimpulsive decisions) than in long-term delays (Gui et al., 2016). Individuals who exhibit larger P3 amplitudes are regarded as being more impulsive, having less mature cognitive control and engaging in more risky (Li et al., 2012; Polezzi et al., 2010). Children’s preference for immediate rewards may be due to their immature cognitive control and their willingness to engage in high-risk behavior; more evidence is needed to investigate the relationships among children’s P3 responses, cognitive control and delay discounting.

Reward amount and time delay are two basic factors in delay discounting, and they were found to have dissociable neural activities: mesial prefrontal cortical activity was positively correlated with future reward magnitude, and DLPFC and posterior parietal cortical activity (PPC) were negatively correlated with future reward delay (Ballard and Knutson, 2009). Scheres et al. (2006) discovered that the choice of children aged between 6 and 11 years old was driven by reward immediacy, while for adolescents aged between 12 and 17 years old, their choices were more influenced by reward amount, which may lead to an increased preference for delayed choices with increasing age. However, the findings of how reward amount and time delay affected children’s neural dynamic processes of delay discounting are less well known and require further exploration.

The main aim of the current study was to investigate the neural dynamic differences between children and adults in delay discounting processes. Since children’s experiences with money and time are limited, we adopted a potential real reward design, which was regarded as suitable for children to make every choice as if it was real (Prencipe et al., 2011; Scheres et al., 2013). We designed four choice conditions: small amount and short delay, small amount and long delay, large amount and short delay and large amount and long delay; we further investigated whether varied reward amounts and time delays might influence this developmental neural process. We chose children in middle childhood as participants because younger children may lack of adequate money and time experience and individuals in late childhood may be more cognitively similar with adolescents. Therefore, children in middle childhood was considered to be a more representative age group for children. It was hypothesized that children might be more likely to choose immediate monetary choices than adults. Children may exhibit smaller and delayed frontal P2 and N2 responses than adults, and children’s developmental lags may be due to their low efficiency in the early detection of rewards in their immature frontal areas. For P3 responses, more neural efforts were needed in children to complete the task by inducing larger P3 responses (Jonkman et al., 2003). In addition, a tendency for risk-taking may positively influence P3 amplitudes (Polezzi et al., 2010), and children might be more likely to engage in risky behavior than adults and thus prefer immediate rewards. Therefore, we further speculated that P3 responses would be larger in children than in adults.

## Results

### Behavioral results

To analyze decision-making processes among varied reward amounts and delays in pre-adolescent children and adults (see Figure 1), the k values, ln(k), RTs and ratios of immediate choices are calculated. The means and standard errors of these variables are presented in Table 1. The different ratios of delayed reward choices of children and adults for varied delays of time are displayed in Figure 2. The ANOVA results on the RTs and the ratios of immediate choices are shown in Tables 2 and 3, and the ratios of immediate choices in different experimental conditions are displayed in Figure 3.

**Table 1.** Means and standard errors (SEs) of k, ln(k), RT and the ratio of immediate choices in children and adults.

| | | RT | $k$(SE) | $\ln(k)$(SE) | Ratio (SE) |
| --- | --- | --- | --- | --- | --- |
| **children** | female | 2295(133) | 0.28(0.07) | -2.35(0.38) | 0.54(0.05) |
|  | male | 1985(87) | 0.33(0.09) | -2.40(0.33) | 0.55(0.04) |
|  | total | 2113(77) | 0.31 (0.06) | -2.38(0.25) | 0.55(0.03) |
| **adult** | female | 1789(176) | 0.01 (0.003) | -4.86(0.19) | 0.23 (0.02) |
|  | male | 1978(149) | 0.02 (0.01) | -4.43 (0.24) | 0.32 (0.03) |
|  | total | 1887(114) | 0.02(0.004) | -4.64(0.16) | 0.28(0.02) |

**Table 2.** Main and interaction effects in the ANOVA analyses for reaction time.

|  | **F** | **P** | **$\eta^2$** |
| --- | --- | --- | --- |
| **Age** | 3.98 | **<u>0.049</u>\*** | 0.04 |
| **Gender** | 0.13 | 0.72 | 0.001 |
| **Age× Gender** | 3.37 | 0.07 | 0.03 |
| **Reward amount** | 9.47 | **<u>0.003</u>\*\*** | 0.09 |
| **Reward amount× Age** | 7.90 | **<u>0.006</u>\*\*** | 0.07 |
| **Reward amount× Gender** | 0.84 | 0.36 | 0.01 |
| **Time delay** | 11.19 | **<u>0.001</u>\*\*** | 0.10 |
| **Time delay× Age** | 1.41 | 0.24 | 0.01 |
| **Time delay× Gender** | 2.01 | 0.16 | 0.02 |
| **Reward amount× Time delay** | 11.18 | **<u>0.001</u>\*\*** | 0.10 |

Note: \*Significance $\leq 0.05$ , \*\*Significance $\leq 0.01$ .

**Table 3.** Main and interaction effects in the ANOVA analyses for the ratio of immediate choices.

|  | **F** | **P** | **$\eta^2$** |
| --- | --- | --- | --- |
| **Age** | 46.57 | **<u>< 0.001</u>\*\*** | 0.31 |
| **Gender** | 1.03 | 0.31 | 0.01 |
| **Age× Gender** | 0.93 | 0.34 | 0.01 |
| **Reward amount** | 660.44 | **<u>< 0.001</u>\*\*** | 0.87 |
| **Reward amount× Age** | 4.09 | **<u>0.046</u>\*** | 0.04 |
| **Reward amount× Gender** | 0.07 | 0.79 | 0.001 |
| **Time delay** | 151.10 | **<u>< 0.001</u>\*\*** | 0.60 |
| **Time delay× Age** | 0.03 | 0.86 | 0.00 |
| **Time delay× Gender** | 4.51 | **<u>0.036</u>\*** | 0.04 |
| **Reward amount× Time delay** | 25.86 | **<u>< 0.001</u>\*\*** | 0.20 |
| **Reward amount× Time delay × Age** | 47.46 | **<u>< 0.001</u>\*\*** | 0.32 |

Note: \*Significance $\leq 0.05$ , \*\*Significance $\leq 0.01$ .

**Figure 1.**
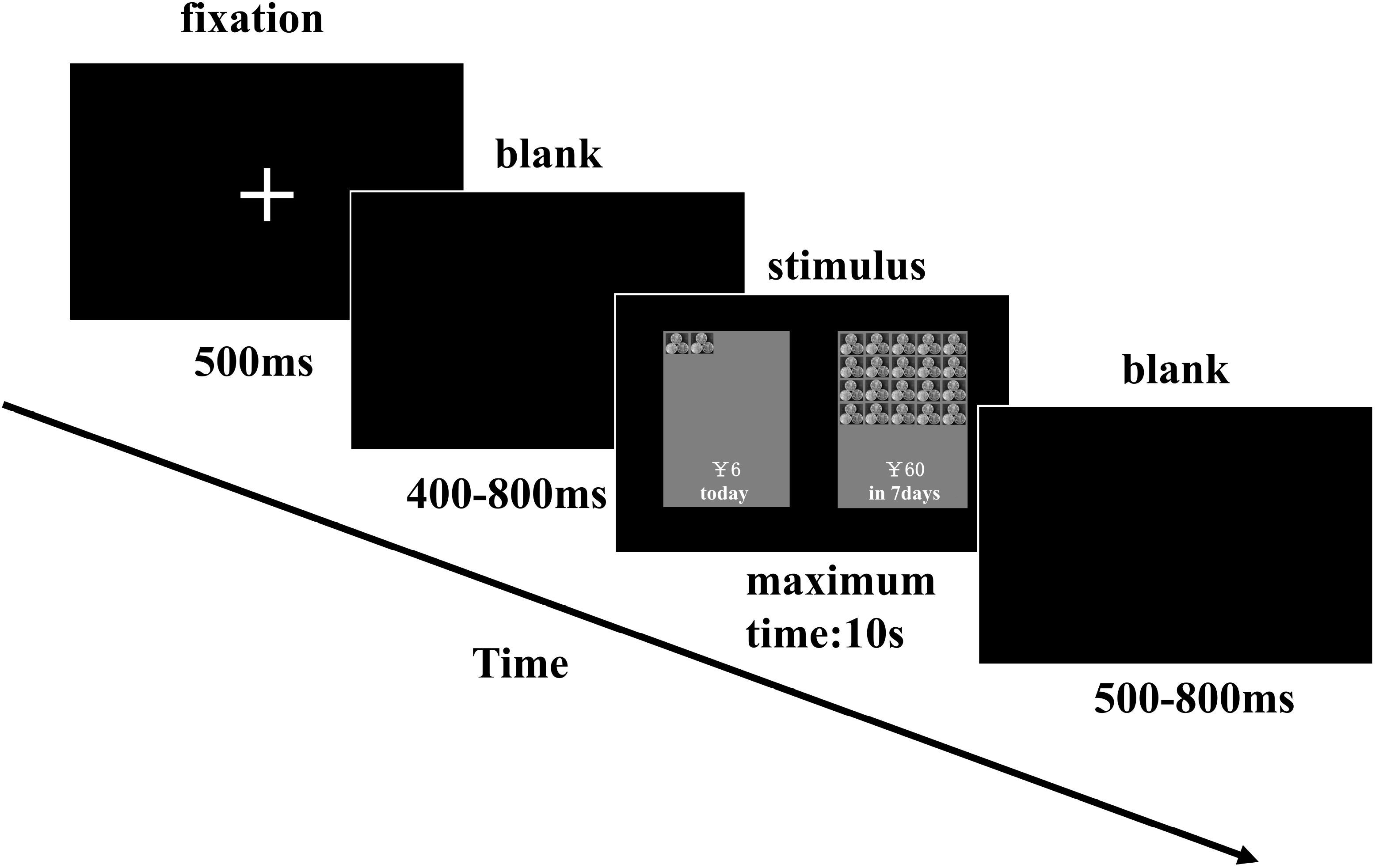
The procedure of the delay discounting tasks.

**Figure 2.**
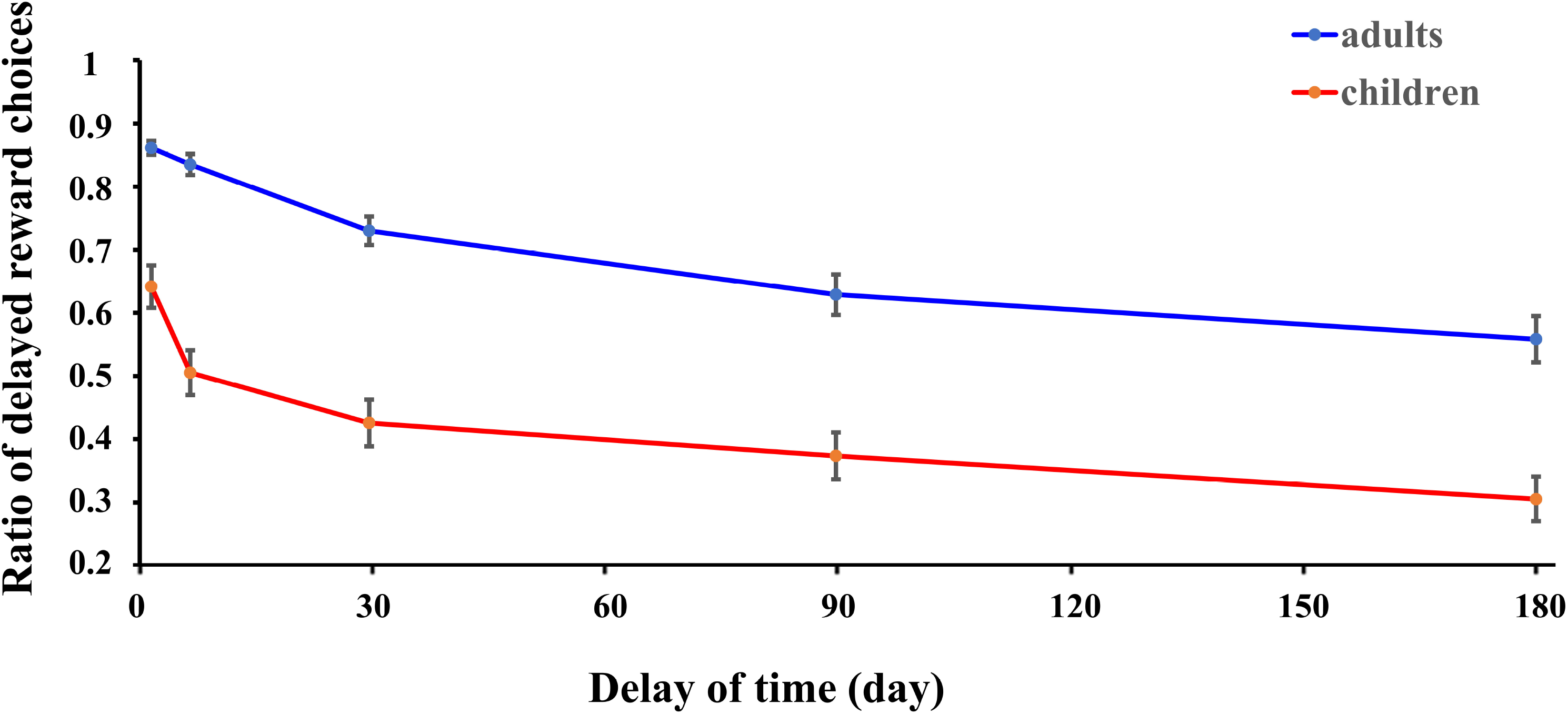
A comparison between children and adults on the ratio of delayed reward choices for various time delays.

**Figure 3.**
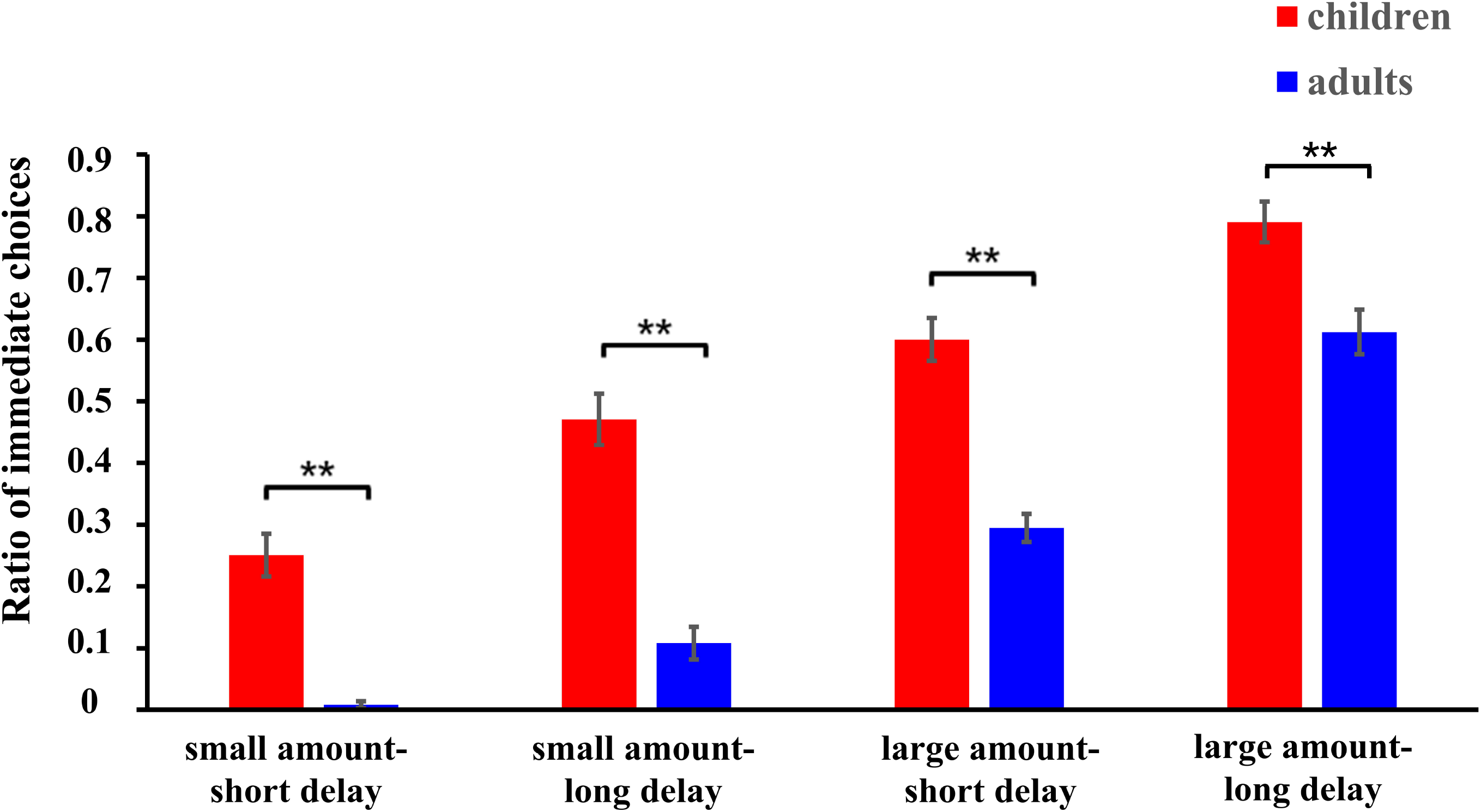
The mean percentage for immediate reward choices during the four conditions in children and adults.

Regarding repeated measures analysis of RTs, a main effect of age was significant, *F*(1, 102) =3.98, *p* = 0.049, *η*^*2*^ = 0.04, in which the RTs of the children were longer than those of the adults. The main effect of reward amount was also significant, *F*(1, 102) =9.47, *p* = 0.003, *η*^*2*^ = 0.09, in which the RT of the large amount was longer than that of the small amount. The main effect of time delay was also significant, *F*(1, 102) =11.19, *p* = 0.001, *η*^*2*^ = 0.10, in which the RTs of short time delays were longer than those of long time delays.

The interaction effect between age and reward amount was significant, *F*(1, 102) =7.90, *p* = 0.006, *η*^*2*^ = 0.07. After the simple-effect tests, it was found that children had longer RTs in the large amount reward condition than in the small amount reward condition (*p* < 0.001), while adults showed similar RTs in both the large and small amount conditions (*p* = 0.86 > 0.05). It was also found that children had significantly longer RTs than adults in the large amount condition (mean _children_ = 2227.12, mean _adults_ = 1890.05, *p* = 0.018) but not in the small amount condition (mean _children_ = 2081.27, mean _adults_ = 1883.48, *p* = 0.14). The interaction effect between reward amount and time delay was also significant, *F*(1, 102) =11.18, *p* = 0.001, *η*^*2*^ = 0.10, and it was observed that in the small amount condition, the RTs of a short time delay were longer than those of a long time delay (mean _short delay_ = 2032.95, mean _long delay_ = 1931.79, *p* < 0.001), while in the large amount condition, there was no significant difference between the RTs of short and long time delays (mean _short delay_ = 2067.77, mean _long delay_ = 2049.40, *p* = 0.40). Additionally, for long time delays, the RTs in the large amount condition were longer than those in the small amount condition (*p* < 0.001), while there were no differences between these two conditions for short time delays (*p* = 0.22)

For the univariate ANOVA of ln(k) values, only the main effect of age was significant, *F*(1, 102) =54.04, *p* < 0.001, *η*^*2*^ = 0.35, in which the ln(k) values of children were larger than those of adults (mean _children_ = −2.37, mean _adults_ = −4.64). There were no other main or interaction effects on ln(k) values.

For the repeated measures analysis of the ratio of immediate choices, the main effect of age was significant, *F*(1, 102) =46.57, *p* < 0.001, *η*^*2*^ = 0.31, in which children chose more immediate choices than adults (mean _children_ = 0.53, mean _adults_ = 0.25). The main effect of reward amount was also significant, *F*(1, 102) =660.44, *p* < 0.001, *η*^*2*^ = 0.87, in which the ratio of immediate choices in the large amount condition was larger than that in the small amount condition (mean _small amount_ = 0.21, mean _large amount_ = 0.57). The main effect of time delay was also significant, *F*(1, 102) =151.10, *p* < 0.001, *η*^*2*^ = 0.60, in which the ratio of immediate choices in the long time delay condition was larger than that in the short time delay condition (mean _short delay_ = 0.29, mean _long delay_ = 0.49).

Furthermore, Spearman correlation analysis showed that the correlation between ln(k) values and the ratio of immediate choice was significant for children (r (58) = 0.98, *p* < 0.001) and adults (r (48) = 0.95, p < 0.001). The larger the ln(k) values was, the more immediate choices individuals would make. The significant results showed consistency across the different analytical methods.

### ERP results

The means and standard errors (SEs) of the latencies and amplitudes of P2, N2 and P3 components are shown in Table 4. We presented only the age-related ANOVA results and hypothesis-relevant post-hoc analyses in the text; all ANOVA results for the amplitudes and latencies are shown in Tables 5 and 6. The grand average waveforms of these maximum amplitudes for ERP components are displayed in Figure 4, and the topographic maps are presented in Figure 5.

**Table 4.** Mean and standard error of ERP latencies and amplitudes in children and adults.

|  |  | **P2** |  | **N2** |  | **P3** |  |
| --- | --- | --- | --- | --- | --- | --- | --- |
|  |  | latency | amplitude | latency | amplitude | latency | amplitude |
| **Children** | SS | 198.41(3.02) | 0.29(0.40) | 245.45(3.58) | -0.19(0.43) | 354.36(3.02) | 6.04(0.44) |
|  | SL | 201.58(2.85) | -0.49(0.37) | 244.78(3.54) | -0.71(0.34) | 356.30(2.76) | 4.78(0.34) |
|  | LS | 197.27(3.12) | -0.44(0.37) | 249.35(3.51) | -1.02(0.42) | 356.72(2.50) | 5.48(0.42) |
|  | LL | 200.41(3.21) | -0.06(0.40) | 237.31(3.72) | -0.20(0.42) | 354.13(2.35) | 5.67(0.35) |
| **Adults** | SS | 178.52(2.83) | 0.44(0.32) | 250.64(3.91) | -1.60(0.33) | 368.94(3.88) | 3.17(0.33) |
|  | SL | 181.64(3.02) | 0.54(0.27) | 250.38(3.78) | -1.40(0.36) | 368.14(3.69) | 3.11(0.29) |
|  | LS | 174.99(2.95) | 0.35(0.27) | 249.76(3.71) | -2.00(0.39) | 367.74(4.00) | 3.19(0.36) |
|  | LL | 178.38(3.00) | 0.48(0.30) | 248.35(3.68) | -1.67(0.42) | 369.33(3.70) | 2.98(0.33) |

Note: SS = small amount and short delay condition; SL = small amount and long delay condition; LS = large amount and short delay condition; LL = large amount and long delay condition

**Table 5.** Main and interaction effects in ANOVA analyses for ERP amplitudes.

| Factors | P2<br>amplitude |  |  | N2<br>amplitude |  |  | P3<br>amplitude |  |  |
| --- | --- | --- | --- | --- | --- | --- | --- | --- | --- |
| | F | p | $\eta^2$ | F | p | $\eta^2$ | F | p | $\eta^2$ |
| Age | 2.16 | 0.14 | 0.02 | 5.06 | **<u>0.027\*</u>** | 0.05 | 28.77 | **<u><0.001\*\*</u>** | 0.22 |
| Gender | 1.19 | 0.28 | 0.01 | 0.19 | 0.67 | 0.002 | 2.68 | 0.111 | 0.03 |
| Reward amount | 0.36 | 0.55 | 0.003 | 1.55 | 0.22 | 0.02 | 0.22 | 0.64 | 0.002 |
| Time delay | 0.12 | 0.73 | 0.001 | 1.43 | 0.24 | 0.01 | 4.41 | **<u>0.038\*</u>** | 0.04 |
| Hemisphere | 20.48 | **<u><0.001\*\*</u>** | 0.17 | 54.37 | **<u><0.001\*\*</u>** | 0.35 | 5.51 | **<u>0.005\*\*</u>** | 0.05 |
| Age × Gender | 0.37 | 0.55 | 0.004 | 0.48 | 0.49 | 0.005 | 0.01 | 0.93 | 0.00 |
| Age × Hemisphere | 4.74 | **<u>0.010\*</u>** | 0.04 | 3.56 | **<u>0.03\*</u>** | 0.03 | 4.02 | **<u>0.019\*</u>** | 0.04 |
| Reward amount × Age | 0.00 | 0.99 | 0.00 | 0.59 | 0.45 | 0.01 | 0.79 | 0.38 | 0.01 |
| Reward amount × Gender | 0.03 | 0.87 | 0.00 | 0.42 | 0.52 | 0.004 | 0.002 | 0.97 | 0.00 |
| Reward amount × Time delay | 4.67 | **<u>0.033\*</u>** | 0.04 | 4.67 | **<u>0.033\*</u>** | 0.04 | 4.21 | **<u>0.043\*</u>** | 0.04 |
| Time delay × Age | 0.91 | 0.34 | 0.01 | 0.41 | 0.52 | 0.004 | 1.74 | 0.19 | 0.02 |
| Time delay × Gender | 0.45 | 0.50 | 0.004 | 2.13 | 0.15 | 0.02 | 0.19 | 0.66 | 0.002 |
| Age × Reward amount × Gender | 4.61 | **<u>0.034\*</u>** | 0.04 | 3.84 | 0.05 | 0.04 | 2.94 | 0.09 | 0.03 |
| Age × Time delay × Gender | 0.01 | 0.92 | 0.00 | 2.13 | 0.15 | 0.02 | 0.72 | 0.40 | 0.01 |
| Age × Time delay × Reward amount | 4.12 | **<u>0.045\*</u>** | 0.04 | 2.86 | 0.09 | 0.03 | 6.29 | 0.01 | 0.06 |

Note: \*Significance $\leq 0.05$ , \*\*Significance $\leq 0.01$ .

**Table 6.** Main and interaction effects in the ANOVA analyses for ERP latencies.

| Factors | P2 latencies |  |  | N2 latencies |  |  | P3 latencies |  |  |
| --- | --- | --- | --- | --- | --- | --- | --- | --- | --- |
| | F | p | $\eta^2$ | F | p | $\eta^2$ | F | p | $\eta^2$ |
| Age | 32.86 | **<u><0.001\*\*</u>** | 0.24 | 1.71 | 0.19 | 0.02 | 11.67 | **<u>0.001\*\*</u>** | 0.10 |
| Gender | 0.100 | 0.75 | 0.001 | 1.50 | 0.22 | 0.01 | 0.34 | 0.56 | 0.003 |
| Reward amount | 3.14 | 0.08 | 0.03 | 1.34 | 0.25 | 0.01 | 0.002 | 0.97 | 0.00 |
| Time delay | 3.95 | **<u>0.049\*</u>** | 0.04 | 3.91 | 0.05 | 0.04 | 0.001 | 0.97 | 0.00 |
| Hemisphere | 7.92 | **<u><0.001\*\*</u>** | 0.07 | 6.66 | **<u>0.002\*\*</u>** | 0.06 | 3.89 | **<u>0.02\*</u>** | 0.04 |
| Age × Gender | 0.04 | 0.84 | 0.00 | 1.67 | 0.20 | 0.02 | 1.13 | 0.29 | 0.01 |
| Age × Hemisphere | 1.76 | 0.17 | 0.02 | 2.20 | 0.11 | 0.02 | 5.82 | **<u>0.003\*\*</u>** | 0.05 |
| Reward amount × Age | 1.07 | 0.30 | 0.01 | 0.002 | 0.97 | 0.00 | 0.03 | 0.86 | 0.00 |
| Reward amount × Gender | 0.07 | 0.80 | 0.001 | 0.47 | 0.50 | 0.01 | 1.40 | 0.24 | 0.01 |
| Reward amount × Time delay | 0.01 | 0.91 | 0.00 | 2.34 | 0.13 | 0.02 | 0.05 | 0.83 | 0.00 |
| Time delay × Age | 0.03 | 0.86 | 0.00 | 2.36 | 0.13 | 0.02 | 0.01 | 0.93 | 0.00 |
| Time delay × Gender | 0.96 | 0.33 | 0.01 | 2.59 | 0.11 | 0.03 | 5.17 | **<u>0.025\*</u>** | 0.05 |
| Age × Reward amount × Gender | 1.92 | 0.17 | 0.02 | 1.35 | 0.25 | 0.01 | 4.50 | **<u>0.036\*</u>** | 0.04 |
| Age × Time delay × Gender | 0.58 | 0.45 | 0.01 | 0.03 | 0.86 | 0.00 | 8.13 | **<u>0.005\*\*</u>** | 0.07 |
| Age × Time delay × Reward amount | 0.003 | 0.96 | 0.00 | 1.62 | 0.21 | 0.02 | 1.05 | 0.31 | 0.01 |

Note: \*Significance $\leq 0.05$ , \*\*Significance $\leq 0.01$ .

**Figure 4.**
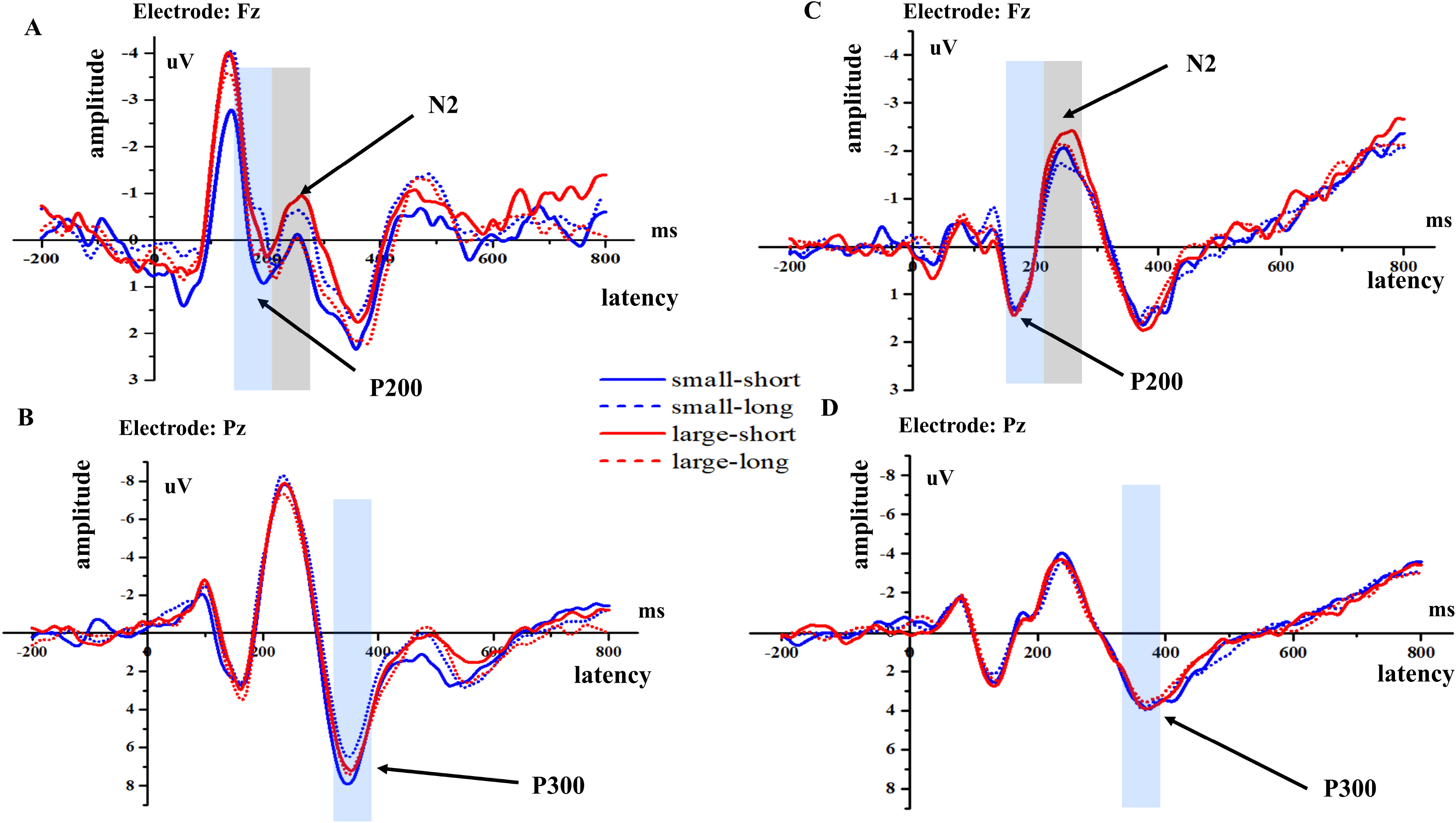
The grand-averaged N2 and P2 ERP waveforms at electrode Fz (Figure 4A) and P3 waveforms at electrode Pz (Figure 4B) in children. Figure 4C & 4D shows the N2 and P2 waveforms at Fz and P3 waveforms at Pz in adults.

**Figure 5.**
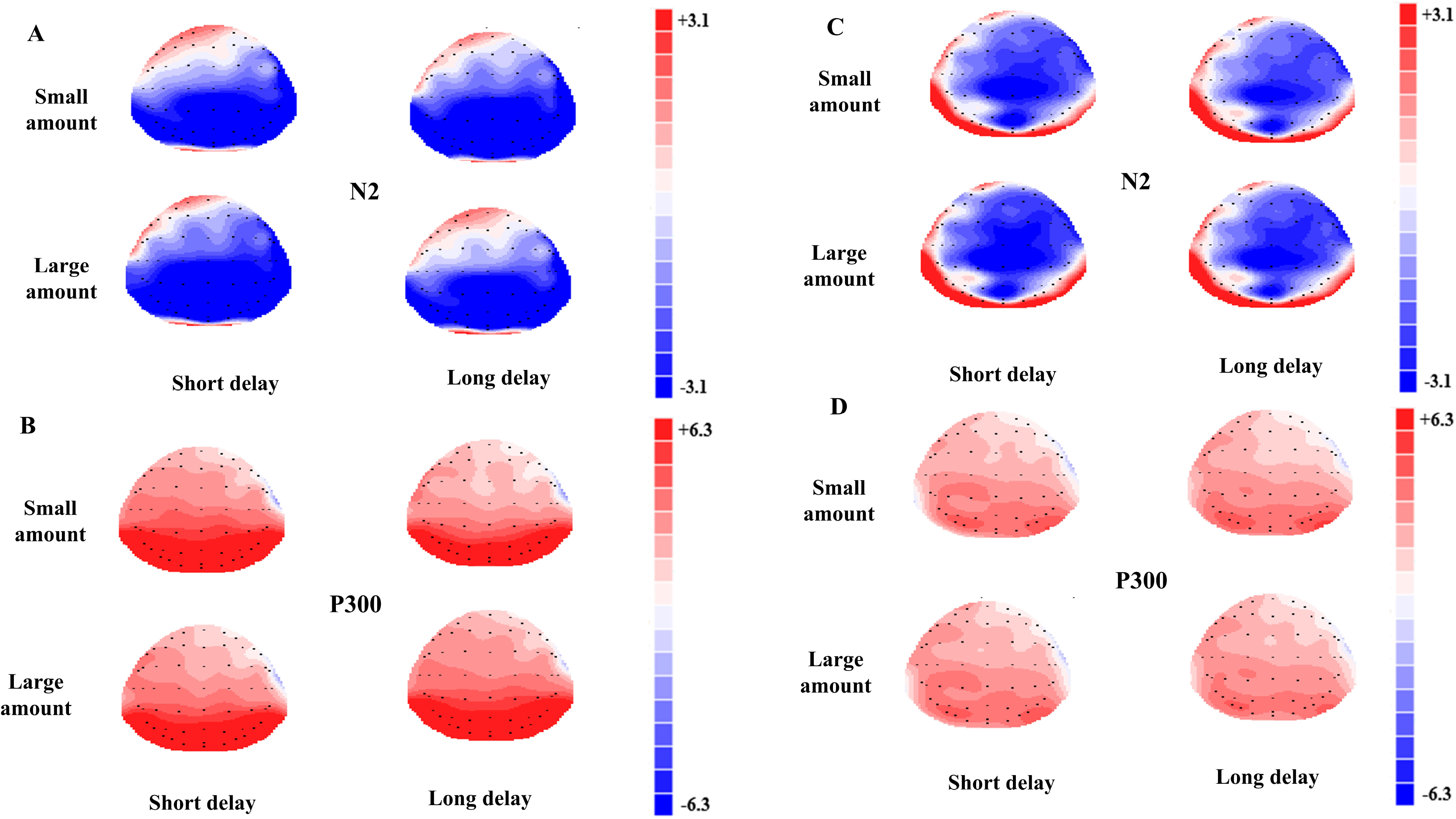
The topographic maps for N2 (Figure 5A) and P3 (Figure 5B) components in children. Figure 5C and 5D shows the topographic maps for N2 and P3 components in adults.

### The P2 component

With regard to P2 amplitudes, the interaction effect of age and hemisphere was significant, *F*(2, 204) =4.74, *p* = 0.010, *η*^*2*^ = 0.04. After the simple-effect test, the amplitudes of children on the right side were significantly smaller than those of adults (*p* = 0.038), while the amplitudes of children were statistically comparable to those of adults in the left and middle brain areas (left: *p* = 0.09; middle: *p* = 0.66).

The interaction effect of reward amount, time delay and age was also significant, *F*(1, 102) =4.12, *p* = 0.045, *η*^*2*^ = 0.04. In the small amount and long delay condition, P2 amplitudes of children were smaller than those of adults (mean _children_ = −0.52, mean _adults_ = 0.56, *p* = 0.025), while in the small amount and short delay condition, in large amount and short delay condition and in large amount and long delay condition, P2 amplitudes of children were not different from those of adults (*ps* > 0.05). In addition, in the small amount conditions for children, the P2 amplitudes in the short delay conditions were larger than those in the long delay conditions (*p* = 0.01), while there were no differences between P2 amplitudes in the short delay and long delay conditions for children in the large amount condition or for adults in both the large and small amount conditions (*ps* > 0.05). Moreover, for children in the short delay conditions, the P2 amplitudes of in the small amount conditions were larger than those in the large amount conditions (*p* = 0.016), but no significant differences were observed between the small and large amount conditions for children in the long delay conditions or for adults in both the short and long delay conditions (*ps* > 0.05).

With regard to P2 latencies, the main effect of age was significant, *F*(1, 102) =32.86, *p* < 0.001, *η*^*2*^ = 0.24, in which children had longer P2 latencies than adults (mean _children_ = 199.58, mean _adults_ = 178.39).

### The N2 component

With regard to N2 amplitudes, the main effect of age was significant, *F*(1, 102) =5.06, *p* = 0.027, *η*^*2*^ = 0.05, in which the amplitudes of children were smaller than those of adults (mean _children_ = −0.54, mean _adults_ = −1.66). The interaction effect of age and hemisphere was significant, *F*(2, 204) =3.56, *p* = 0.030, *η*^*2*^ = 0.03. After a simple-effect test, the N2 amplitudes of children were significantly smaller than adults over the middle and right areas (*p* (right) = 0.021; *p* (middle) = 0.007), while the activities of both age groups were not significantly different on the left brain areas (*p* (left) = 0.16).

The interaction of reward amount, age and gender was marginally significant, *F*(1, 102) = 3.84, *p* = 0.05, *η*^*2*^ = 0.04. It was further found that in the small amount conditions for males, the N2 amplitudes of children were smaller than those of adults (mean _children_ = −0.21, mean _adults_ = −1.88, *p* = 0.012), while no differences were found between the amplitudes of children and adults in the small amount conditions for females or in the large amount conditions for males and females (small amount, female: mean _children_ = −0.79, mean _adults_ = −1.09, *p* = 0.68; large amount, female: mean _children_ = −0.43, mean _adults_ = −1.67, *p* = 0.14; large amount, male: mean _children_ = −0.74, mean _adults_ = −2.00, *p* = 0.10).

With regard to N2 latencies, there were no age-related main or interaction effects on N2 latencies.

### The P3 component

With regard to P3 amplitudes, the main effect of age was significant, *F*(1, 102) =28.77, *p* < 0.001, *η*^*2*^ = 0.22, in which children had larger P3 amplitudes than adults (mean _children_ = 5.55, mean _adults_ = 3.13). The interaction effect of reward amount, time delay and age was significant, *F*(1, 102) =6.29, *p* = 0.014, *η*^*2*^ = 0.06. After the simple effect test, in small amount condition, children had larger P3 amplitudes for the short delay conditions than for the long delay conditions (children, small amount: mean _short_ = 6.06, mean _long_ = 4.83, *p* < 0.001), while in the large amount conditions for children and in small and large amount conditions for adults, P3 amplitudes in the short delay conditions were not different from those in the long delay conditions (children, large amount: mean short = 5.58, mean _long_ = 5.73, *p*=0.62; adults, small amount: mean _short_ = 3.19, mean long = 3.14, *p* = 0.87; adults, large amount: mean _short_ = 3.2, mean _long_ = 3.00, *p* = 0.55). In addition, for children in the long delay conditions, the amplitudes of small amounts were smaller than those of large amount (*p* = 0.002), while for children in the short delay conditions and for adults in the short and long delay conditions, there were no significant differences between the amplitudes of small amounts and large amounts (children, short delay, small amount & large amount: *p* = 0.12; adults, short delay, small amount & large amount: *p* = 0.99; adults, long delay, small amount & large amount: *p* = 0.66).

With regard to P3 latencies, the main effect of age was significant, *F*(1, 102) =11.67, *p* = 0.001, *η*^*2*^ = 0.10, in which children had shorter P3 latencies than adults (mean _children_ = 355.9, mean _adults_ = 368.5, *p* = 0.001). The interaction effect of reward amount, gender and age was significant, *F*(1, 102) =4.503, *p* = 0.036, *η*^*2*^ = 0.04. A simple effect test revealed that in both the small and large amount conditions for males, the latencies of children were significantly longer than those of adults (small amount, male: mean _children_ = 366.99, mean _adults_ = 353.41, *p* = 0.02; large amount, male: mean _children_ = 371.77, mean _adults_ = 352.32, *p* < 0.001); there were no other significant differences for females (small amount, female: mean _children_ = 370.23, mean _adults_ = 358.05, *p* = 0.06; large amount, female: mean children = 365.01, mean _adults_ = 359.81, *p*=0.34).

The interaction effect of time delay, gender and age was significant, *F*(1, 102) = 8.13, *p* = 0.005, *η*^*2*^ = 0.07. A simple effect test revealed that for female children, the P3 latencies in the short delay conditions were longer than those in the long delay conditions; for male children, the P3 latencies in the short delay conditions were shorter than those in the long delay conditions, while for female adults and male adults, the latencies in the short delay and long delay conditions did not differ (children, female: mean _short_ = 371.45, mean _long_ = 363.79, *p* = 0.019; children, male: mean _short_ = 365.48, mean _long_ = 373.29, *p* = 0.013; adults, female: mean _short_ = 358.58, mean _long_ = 359.28, *p* = 0.82; adults, male: mean _short_ = 353.39, mean _long_ = 352.34, *p* = 0.69). In addition, it was also observed that for females in the short delay conditions, for males in the short delay conditions and for males in the long delay conditions, the latencies of children were longer than those of adults (female, short delay, *p* = 0.034; male, short delay, *p* = 0.028; male, long delay, *p* < 0.001), while for females in the long delay conditions, the latencies of children were not different from those of adults (female, long delay, *p* = 0.44).

### Correlation between behavioral and ERP data

The correlations between the ratio of immediate choices and ERP responses and that between the ln(k) values and ERP responses are presented in Table 7. There were significant correlations between P3 amplitudes and ln(k) values (*r* (106) = 0.37, *p* < 0.001), between P3 amplitudes and the ratio of immediate choices (*r* (106) = 0.31, *p* = 0.001) and between P3 latencies and ln(k) values (*r* (106) = −0.23, *p* = 0.018), and between P3 latencies and the ratio of immediate choices (*r* (106) = −0.21, *p* = 0.033). Moreover, there were significant differences between P2 latencies and ln(k) values (*r* (106) = 0.30, *p* = 0.002) and between P2 latencies and the ratio of immediate choices (*r* (106) = 0.32, *p* = 0.001).

**Table 7.**
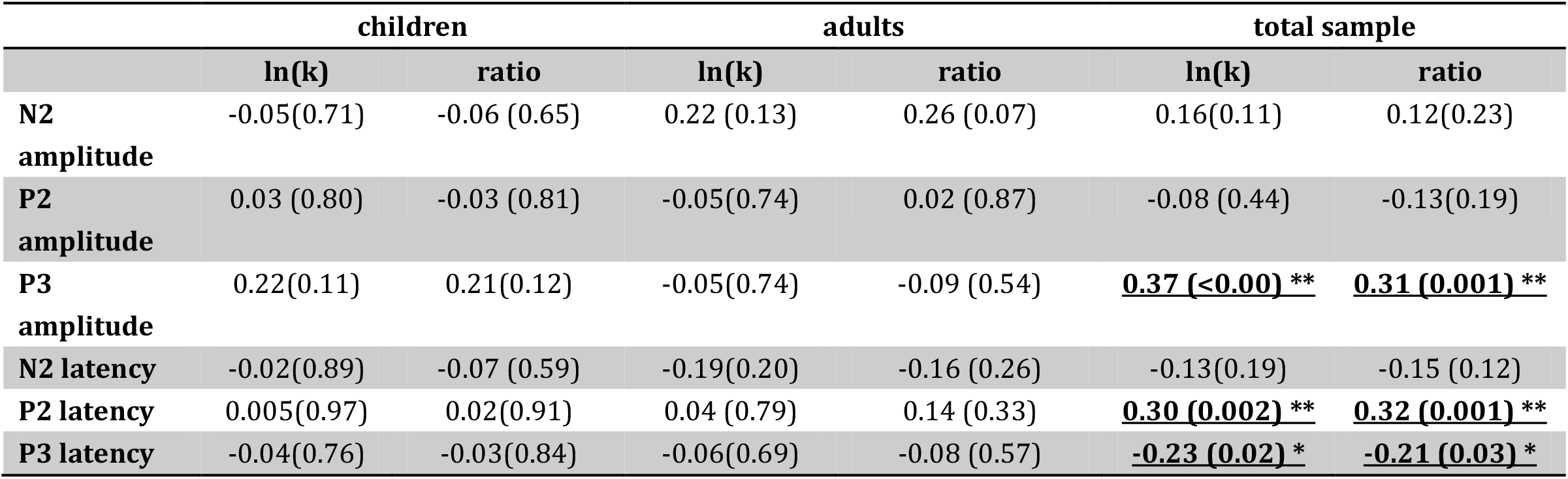
Pearson’s correlations among ln(k) values, ERP component amplitudes and ERP component latencies, and Spearman’s correlations among the ratio of immediate choices, ERP component amplitudes and ERP component latencies.

## Discussion

The goal of the study was to discover the temporal dynamics underlying the behavioral differences in delay discounting processes of children and adults. The current study showed that children discounted more than adults in behavioral performance and exhibited longer frontal P2, smaller N2 and larger P3 amplitudes than adults. In addition, we found that reward amount and time delay influenced children and adults differently.

### Age differences in delay discounting

The behavioral findings showed that children’s discount rate was higher than that of adults and that children made more immediate choices than adults, which was consistent with previous findings (Green et al., 1994; Olson et al., 2007; Steinberg, 2009). Valuation and cognitive control are very important for delay discounting (Peters and Buchel, 2011), and these findings regarding developmental differences might be because children have less mature valuation and cognitive control functions than adults. Compared to adults, children showed higher sensitivity to rewards (Galvan et al., 2006), and they had worse cognitive control function than adults, including working memory and response inhibition (Huizinga et al., 2006; Luciana et al., 2005). Moreover, children may lack experience with long delays and may view the same range of time as longer than adults perceive it to be (Green et al., 1994; Olson et al., 2007), which may affect their future orientation. Steinberg et al. found that younger individuals who showed a higher discount rate exhibited a weaker orientation to the future, especially in the dimensions of temporal orientation and the anticipation of future consequences but not in the dimension of planning ahead (Steinberg, 2009). In fact, children’s concept of the future is formed during development (Silverman, 1996). In addition, different aspects of future orientation might have different developing trajectories since different dimensions of future orientation have different neural bases (Fellows and Farah, 2005), which may lead to different influences on delay discounting. Regardless, it is likely that children experience long-time delays less than adults, which makes children less future oriented and more likely to choose immediate choices. Overall, children’s steeper discounting may result mainly from aspects of their cognitive ability and experience, but how the two aspects interact needs to be further investigated in the future.

Neuroimaging studies have found that the activation of the limbic system associated with the midbrain dopamine system is involved in immediately available rewards, while greater relative frontoparietal activation is related to longer-term options (McClure et al., 2004). In children, the “bottom-up” brain regions related to reward valuation, such as the ventral striatum (VS), develop faster than the “top-down” brain cortexes involved in cognitive control, such as the DLPFC and parietal cortex (Blakemore and Choudhury, 2006; Giedd et al., 1999; Gogtay et al., 2004; Scheres et al., 2013; Stanger et al., 2013). Notably, children had significantly longer RTs than adults for the large amount rewards but not for the small amount rewards, which revealed that it took children more time to decide on large amount immediate rewards than on small amount immediate rewards.

The P2 component is related to early feature encoding on stimuli and attention filtering in the early decision-making period (Harris et al., 2013; Li et al., 2012; Potts, 2004), and the present study observed that children had longer P2 responses than adults. A previous study found that high procrastinators exhibited longer P2 responses in delay discounting, which showed abnormal reward encoding (Wu et al., 2016). This finding indicated that children might be slower during early encoding in delay discounting. Moreover, children showed smaller P2 amplitudes than adults only in the small amount and long delay condition; thus, the features of small amounts and long delays may attract children less than adults during early processing periods.

During delay discounting, the brain areas involved in cognitive control were cortexes such as the lateral prefrontal cortex (McClure et al., 2004; Peters and Buchel, 2011). The frontal N2 componnet is regarded as a neural marker for individual differences in executive functions such as cognitive control (Espinet et al., 2012). Individuals suffering from earthquakes have been shown to have smaller N2 responses and to have a higher degree of discounting (Li et al., 2012); also, their N2 amplitude was negatively correlated with the ratio of immediate choices (Gui et al., 2016). Our current study found that children had smaller N2 amplitudes than adults over the middle and right frontal areas, which might indicate children’s immature development in cognitive control and is consistent with the slower development of the top-down processing system in decision making (Blakemore and Choudhury, 2006; Giedd et al., 1999; Gogtay et al., 2004). This finding was also in agreement with the fMRI finding that children’s ability to overcome temptation increased with improved in functional coupling between the VMPFC, which is related to reward valuation, and brain regions such as the DLPFC, which are involved in behavioral control (Steinbeis et al., 2016). In addition, the current age-related differences were over middle and right frontal areas but not left areas, similar to that reported in other studies. Stanger et al. (2013) found that the activation of right frontal-parietal areas correlated more with ln(k) values than the activation of left frontal-parietal areas, which indicated that individuals exhibiting higher degrees of discounting showed a larger right hemisphere effect. Additionally, the inhibition control process induced stronger activation over the right and middle areas of the frontal lobe, insula and parietal lobe (Garavan et al., 1999). Therefore, it is plausible that the age differences in the delay discounting task, involving cognitive control are exhibited mainly in the right hemisphere. More fMRI studies should be performed to further test this hypothesis.

The P3 component is regarded as a component to investigate various cognitive processes, processing capacity and mental workload (Kok, 2001; Polich, 2007). In the present study, children had larger P3 responses than adults. Male children had longer P3 latencies than male adults, and female children had longer P3 latencies than female adults in short delay conditions. This result could be explained as follows. First, in line with several developmental studies, the findings of the current study suggest that children recruited more neural resources to execute decision-making processes than adults (Jonkman et al., 2003). Casey et al. (1997) observed that children exhibited stronger activation in dorsolateral prefrontal areas than adults when accomplishing a go/nogo task. Moreover, it was discovered that during childhood, working memory is still developing (Huizinga et al., 2006) and that delay discounting is negatively correlated with working memory (Shamosh et al., 2008). Second, it has been shown that higher motivation could induce greater P3 in delay discounting (Patalano et al., 2018; Polich and Kok, 1995), and previous fMRI studies showed that reward-related brain regions such as the accumbens showed larger activation in children than in adults (Galvan et al., 2006). Hence, children’s larger P3 responses during delay discounting in the current task might be related to their higher sensitivity and motivation to rewards. Third, children might view future rewards as an event with a lower possibility of occurrence since they have less experience with long time delays. From an evolutionary perspective, future rewards have been regarded as uncertain (Fehr, 2002). Furthermore, a previous study revealed that a lower probability of an infrequent target led to increased P3 amplitudes (Carlson et al., 2009). From these findings, it can be concluded that children may consider future rewards to be uncertain, which influences their decision-making process in delay discounting, while adults may be less influenced by the uncertainty of future rewards. Fourth, there were significant interaction effects of age and gender factors on P3 latencies: female children had slower P3 responses than female adults mainly in short delay conditions, while male children showed slower P3 responses in both short and long delay conditions. These findings illustrated that female children’s developmental lag in delay discounting was obvious in the short delay condition, while male children’s lag existed in both delay conditions.

### The influence of reward amount and time delay on the development of delay discounting

In the current study, the ratio of immediate choices was higher in large amount trials than in small amount trials, and it was higher in long delay trials than in short delay trials. According to our study design, a change in the reward amount occurred in the immediate options; hence, while a change in the time delay occurred in the delay options; it is understandable that individuals would make immediate choices more when the immediate amount was large or the delay time was long.

There are some influences of reward amount and time delay on age and gender differences. Children spent a longer time making a choice in the large amount condition than in the small amount condition, while adults had similar RTs in both the small and large amount conditions. For neural dynamic processes, in the small amount condition, children exhibited larger P2 and P3 amplitudes in the short delay condition than in the long delay condition. In the short delay condition, children had larger P2 amplitudes in the small amount condition than in the large amount condition. In the long delay condition, children had smaller P3 responses in the small amount condition than in the large amount condition. However, in all the above situations, adults showed no differences across conditions. Actually, choices in the small amount and short delay condition were easier for children than those in the small amount and long delay condition since the prior choices consisted of one unfavorable option in the amount dimension, while the other option favored the time dimension; the latter choices consisted of one unfavorable option in the amount dimension and the another unfavorable option in the time dimension. It can be concluded that for children, easier choices evoked larger P3 amplitudes. It is possible that children could employ more neural resources to compute or evaluate the reward for easier choices than for more difficult choices. The results may also be explained by the motivational significance of children for the easier choices, as in the evaluation period, P3 amplitudes may also be influenced by motivation (Kok, 2001), with more motivated stimuli evoking larger P3 amplitudes (Schupp, 2003).

Moreover, female children had longer P3 latencies in the short delay conditions than in the long delay conditions, but female children had shorter P3 latencies in the short delay conditions than in the long delay conditions. However, female and male adults did not exhibit any differences between the short and long delay conditions. It is believed that those who more strongly discounted the value of delayed rewards more deeply experience time to have a higher cost and often overestimate the duration of time intervals (Wittmann and Paulus, 2008). A previous study using time reproduction showed that children often overvalue time (Szelag et al., 2002); in other words, children often overestimate the duration of time. In addition, it was discovered that adults’ time sensitivity was higher than children (Droit-Volet et al., 2007); that is, adults made fewer errors and estimated time more correctly than children. Therefore, it seems that children’s lower sensitivity to time delays may lead to higher degrees of discounting. According to this hypothesis, when the time delay increases, children may show a higher degree of overestimation when estimating longer delays than when estimating shorter delays, which may lead to a higher degree of discounting in longer delays than in shorter delays. However, our results contradict this hypothesis, as children exhibited comparable differences to adults in both short and long delays. The reason that children’s long time estimation for 2 or 7 days and for 30, 90 or 180 days in our study was comparable may be because children have less experience in long time delays, and the estimation pattern in long time delays is different from that in short time delays, which are often used in the laboratory.

In conclusion, the current study found that children discounted monetary rewards more steeply than adults and more likely chose immediate choices. Regarding neural dynamic processes, children showed longer early detection and identification processing for the presented choices than adults. They also showed less mature cognitive control processing over frontal areas and devoted more neural efforts to make choices during the decision phase. Moreover, the factors of reward amount and time delay could affect male children’s decision making, with male children showing slower neural responses than male adults in the long delay and large reward amount conditions. The present study sheds light on the neural development of delay discounting in children and adults.

### Materials and methods Participants

The study was approved by the local Ethics Committee of the Institute of Psychology, Chinese Academy of Science and was conducted in accordance with the Declaration of Helsinki. Two age groups participated in the current study. Sixty-two children were recruited from a local elementary school by advertisement. Two children were excluded due to recording errors of the instrument, one child was excluded due to attention problems, and one child was excluded because there were not enough trials suitable for neural analysis. Thus, 58 children (34 boys; 9-10 years old; mean age: 9.26±0.44 years) were included for further analysis. For the adult group, 50 adults were recruited from local universities by advertisement; one adult was excluded due to a different sampling rate, and one adult was excluded due to a history of depression. Therefore, there were 48 adults (25 males; aged 18-26 years old; mean age: 20.98±1.93 years) included for further analysis. Child participants were awarded for participation with a present after completion of the task, and adult participants were paid for participation. All participants were right-handed with normal or corrected-to-normal vision, reported being free from psychiatric problems, and did not regularly use medications. Written informed consent was provided by parents of the children and by adult participants before the experiment.

### Experimental stimuli and paradigm

The current task was an adaption of Mitchell’s paradigm (Mitchell, 1999). The participants were seated comfortably in front of a computer and performed the task in an electromagnetically shielded room. The stimuli were presented on a computer monitor with a 60 Hz refresh rate, and the acquisition of behavioral data was conducted with E-Prime software (Version 2.0, Psychology Software Tools, Inc.).

For each trial, two options were presented on the right and left sides of the screen simultaneously. One was an immediate and smaller monetary reward, and the other was a delayed and larger monetary reward. Participants were asked to choose one of the two options. The immediate options referred to a situation in which participants could gain the monetary reward immediately after the experiment. The amount of immediate monetary rewards changed from ¥0 to ¥63 in the step of ¥3, in which ¥0~¥30 were regarded as the small amount and ¥33~¥63 were regarded as the large amount. The delayed options referred to the situation in which participants could gain a fixed amount of money (¥60) in varied delayed times from 2 days to 180 days (detailed delays: 2, 7, 30, 90 or 180 days), in which 2 and 7 days were deemed short time delays and 30, 90 and 180 days were deemed long time delays. The immediate rewards and delayed times were selected randomly to form the choices. Before the formal experiment, there were 8 practice trials, which were used to make subjects understand the rules of the task. In the formal task, there were 44 trials for the small amount (immediate option) and short delay (delay option) condition, 66 trials for the small amount (immediate option) and long delay (delay option) condition, 44 trials for the large amount (immediate option) and short delay (delay option) condition, and 66 trials for the large amount (immediate option) and long delay (delay option) condition. Participants were required to have a 2-3 minute rest period every 74 trials.

The procedure is presented in Figure 1. Each trial began with the presentation of a fixation point for 500 ms, followed by a blank screen for a randomized duration ranging from 400 to 800 ms. Then, the choice screen was presented with a maximum time of 10 s, which was long enough for all child and adult participants to make a choice. The intertrial interval was a random period of 500 to 800 ms. The participants indicated their choices by pressing ‘z’ with their left index fingers if they preferred the option on the left side of the screen and ‘/’ with their right index fingers if they preferred the option on the right side of the screen. The presentation of the choice pairs and the location of the immediate and delayed choices in each pair were random.

Before the experiment, we informed all the participants about the reward rules. The rewards in the current task were potentially real. Participants were explicitly required to choose the reward options according to their real preference because they would be given one of their choices randomly after the experiment (Scheres et al., 2013). Participants clearly knew that they could receive the monetary reward immediately after finishing the experiment if they chose the immediate option. Participants were instructed that if they chose a delayed reward option, the money would be transferred to their bank account (adult participants) or given to them in an envelope by their parents (child participants) on the exact day that they chose in the selection phase.

### ERP data acquisition

Electroencephalograms (EEGs) were recorded from a Neuroscan Quick-Cap (64 scalp sites) according to the International 10/20 system, with an online reference to the nose. Horizontal electrooculogram (HEOGs) were recorded from electrodes placed approximately 1.5 cm lateral to the left and right external canthi. Vertical electrooculograms (VEOGs) were recorded from the left supraorbital and infraorbital electrodes. All interelectrode impedances were maintained below 10 kΩ. All signals were sampled at 1000 Hz and online-bandpass filtered within a 0.05–100 Hz frequency range.

For off-line analysis, EEG signals affected by body movement were removed from further analyses, and ocular artifacts were corrected with an eye-movement correction algorithm implemented in Neuroscan software (Semlitsch et al., 1986). Trials containing EEG sweeps with amplitudes exceeding ±90 μV were excluded. EEG data were filtered from 1 to 30 Hz (24 dB/oct). EEG signals were divided into epochs of 1000 ms (together with 200 ms prestimulus as the baseline), time locked to the onset of stimulus. The trial numbers left for further statistical analysis in each condition were as follows. For children, there were 37± 5 trials in the small amount and short delay condition, 55 ± 7 trials in the small amount and long delay condition, 35 ± 6 trials in the large amount and short delay condition and 55 ± 8 trials in the large amount and long delay condition. For adults, there were 43 ± 2 trials in the small amount and short delay condition, 64 ± 3 trials in the small amount and long delay condition, 42 ± 2 trials in the large amount and short delay condition, and 64 ± 3 trials in the large amount and long delay condition.

### Data analysis

For behavioral data, three parameters were analyzed: the ratio of immediate choices, the discount rate k (Mazur, 1987; Odum et al., 2006), and the reaction times (RTs) of the choices. The k value was calculated from the hyperbolic model (Mazur, 1987), which is written as:

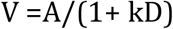

where V shows the subjective value of an outcome. We obtained the k value from the turning point, which was the change from delayed choices to immediate choices noted after arranging the immediate qualities of each delay time. A is the objective value of V. In our study, A was fixed at ¥60. D represents the delay to its receipt. In our study, D was 2, 7, 30, 90 or 180 days. After entering V, A and D in the formula, we obtained a value for k, the rate of discounting. Because the data of k were not normally distributed and therefore not suitable for further analysis, we transformed k to ln(k), as in a previous study (Wilson et al., 2011). Notably, ln function is an increasing function in which larger ln values indicate a higher discounting rate and smaller values demonstrate a lower discounting rate. For ln(k) values, univariate ANOVAs were conducted with age (child vs. adult) and gender (female vs. male) as between-subject factors. For RTs and the ratio of immediate choices, repeated measures analyses were conducted separately with reward amount (small amount vs. large amount) and time delay (short delay vs. long delay) as within-subject factors and age (child vs. adult) and gender (female vs. male) as between-subject factors.

For ERP data, three ERP components were analyzed-P2, N2 and P3 components-and the time windows and the electrode sites for these components were chosen based on previous literature (Lamm et al., 2006; Polich, 2007; Potts, 2004) and visual inspection of the current ERP grand average waveforms. P2 and N2 components were analyzed over the fronto-central areas (Fz, F3, F4, FCz, FC3, and FC4) during the time windows of 140 to 240 ms and 200 to 300 ms, respectively. The P3 component was measured over the centro-parietal electrodes (Pz, P3, P4, CPz, CP3, and CP4) with a time window of 310 ms to 410 ms.

For the amplitudes and latencies of P2, N2 and P3, repeated measures analyses were conducted separately with reward amount (small amount vs. large amount), time delay (short delay vs. long delay) and hemisphere (left vs. middle vs. right) as within-subject factors and age (child vs. adult) and gender (female vs. male) as between-subject factors.

For correlations of behavioral results and ERP data, we performed Pearson’s correlation between ln(k) values and the amplitudes or latencies of the ERP components. Because the ratio of immediate choices was not normally distributed, we performed Spearman’s correlation analyses (nonparametric test) between ln(k) values and the ratio of immediate choices and between the amplitudes or latencies of the ERP components and the ratio of immediate choices.

For all statistical analyses, we used SPSS version 22.0. Significant interactions were analyzed by post hoc simple effects. Partial eta-squared is represented to demonstrate the effect size of the results.

## Acknowledgement

This research was supported by the National Natural Science Foundation of China (Grant No. 31370020) and the CAS Key Laboratory of Behavioral Science, Institute of Psychology, Chinese Academy of Sciences. We express our thanks to all the children and adult subjects for their participation.

